# Approximating Solutions of the Chemical Master Equation using Neural Networks

**DOI:** 10.1101/2022.04.26.489548

**Authors:** Augustinas Sukys, Kaan Öcal, Ramon Grima

## Abstract

The Chemical Master Equation (CME) provides an accurate description of stochastic biochemical reaction networks in well-mixed conditions, but it cannot be solved analytically for most systems of practical interest. While Monte Carlo methods provide a principled means to probe the system dynamics, their high computational cost can render the estimation of molecule number distributions and other numerical tasks infeasible due to the large number of repeated simulations typically required. In this paper we aim to leverage the representational power of neural networks to approximate the solutions of the CME and propose a framework for Neural Estimation of Stochastic Simulations for Inference and Exploration (Nessie). Our approach is based on training a neural network to learn the distributions predicted by the CME from a relatively small number of stochastic simulations, thereby accelerating computationally intensive tasks such as parameter exploration and inference. We show on biologically relevant examples that simple neural networks with one hidden layer are able to capture highly complex distributions across parameter space. We provide a detailed discussion of the neural network implementation and code for easy reproducibility.

## 1 Introduction

The past decades have seen great progress in our understanding of the complex dynamics that underlie noisy cellular processes, both from an experimental and a theoretical perspective. Modern experimental techniques have shown that mRNA and protein levels can vary enormously at the single-cell level, but building detailed quantitative models that take into account the stochasticity of biochemical systems remains a daunting task. Due to the numerous challenges involved in describing the stochastic dynamics of these systems, modeling frequently relies on deterministic and small noise approximations which do not paint an accurate picture in many situations. Such simplified descriptions are often insufficient to describe how biochemical networks function in the presence of molecular noise [1, 2] and do not capture intricate noise-driven phenomena involved in cell fate decision [3, 4] and phenotypic regulation [5].

The most commonly used formalism for modeling biochemical reaction networks in a fully stochastic framework is the Chemical Master Equation (CME) [6], which describes how the probability distribution over states evolves with time. The CME cannot be solved analytically for most biologically relevant cases, and since the state space is typically infinite, numerical solutions of the CME often involve state space truncation methods such as the Finite State Projection (FSP) [7]. However, due to the combinatorial explosion of the state space in the number of species, using the FSP to solve the CME quickly becomes too computationally intensive for most non-trivial systems [8–10]. A wide variety of other approximation methods exist for the CME (see [6] for an overview), but these often trade computational efficiency for accuracy and are generally difficult to apply to complex systems involving many species and interactions.

While solving the CME remains challenging, *sampling* realizations of a system is possible thanks to the Stochastic Simulation Algorithm (SSA) [11]. Many physical quantities such as moments of molecule number distributions can be computed to arbitrary accuracy by repeatedly simulating samples from the system. Nevertheless, the SSA can be prohibitively computationally expensive when many repeated simulations are needed for accurate estimation of these quantities. Since simulations have to be performed anew for all parameters of interest, investigating system properties over time and parameter space with this approach can quickly become intractable. Furthermore, likelihoods are hard to estimate reliably using Monte Carlo methods, rendering likelihood-based inference particularly difficult [12].

Given the difficulties inherent in solving the CME exactly, it is natural to explore whether we could tackle this problem using neural networks, which in recent years have found diverse applications in the physical and biological sciences [13–16]. Their ability to detect patterns and learn complex representations given enough data is particularly useful when combined with simulator-based modeling, where such data can often be generated aplenty. In the context of systems biology, neural network-based approaches have been used to perform parameter inference on deterministic models, accelerate parameter exploration for models described by partial and stochastic differential equations [17], and even learn Markovian approximations to non-Markov models (translating them into the CME framework) [18], amongst other applications.

Moreover, a number of recent studies have investigated the use of neural networks to learn various properties of stochastic biochemical reaction networks modeled using the CME [19–23]. In [19] the authors presented DeepCME, an approach that uses reinforcement learning to estimate summary statistics such as means and variances from stochastic simulations. The model abstraction procedure introduced in [20] employs Mixture Density Networks [24] to learn the transition kernel of the CME, and has been further extended into a framework providing automated neural network architecture search [21]. In the same vein, mixture density networks have been used to directly predict the probability distributions characterizing the dynamics of an SIR-type model [22]. Finally, in [23] the authors demonstrate how Generative Adversarial Networks can be trained to generate full trajectories resembling the output of the SSA.

In this paper we introduce Nessie, a framework for Neural Emulation of Stochastic Simulations for Inference and Exploration, based on using neural networks to learn solutions of the CME from stochastic simulations. Using only a moderate number of simulations of the specified reaction system at different parameter values, we train a neural network to learn the marginal probability distributions predicted by the CME over the whole parameter region of interest. We approximate the target distributions using mixtures of negative binomials, a flexible class of distributions particularly well-suited for this task [25, 26]. Once trained, Nessie becomes a surrogate for the CME that can efficiently and accurately predict the solution of the CME for a wide range of parameters, bypassing the need to use further simulations or expensive approximation techniques to analyze the reaction network in question.

Our work differs from related approaches [19–23] in several regards. Unlike [20, 21] or [23], we do not aim to learn the transition kernels or the distributions of trajectories, i.e. the dynamics of the chemical system in question, but to capture the *marginal distributions* directly. In this sense Nessie is also different from DeepCME [19], which focuses on the task of learning summary statistics such as moments of molecule numbers. The relevant expectation values can be computed directly from the distributions predicted by Nessie, and we verify in our examples that Nessie can predict means and variances to a high degree of accuracy. Although our neural network is a variant of mixture density networks, in contrast to [20, 21] we do not use continuum approximations based on mixtures of Gaussians, thereby avoiding training and numerical stability issues that can arise in this context [20, 21, 27, 28]. As our approach directly targets experimentally observable distributions, it can also be used to perform parameter estimation based on comparing the neural network output with experimental data.

The paper is organized as follows. In Section 2 we summarize the relevant theory on the CME and the basics of artificial neural networks. In Section 3 we describe Nessie, providing an overview of the technical details of our neural network implementation and the workflow we use to predict the marginal probability distributions for a given system. In Section 4 we test our approach on several biologically relevant examples: an autoregulatory feedback loop, a genetic toggle switch involving mRNA and protein dynamics for two genes and a model of the mitogen-activated protein kinase (MAPK) pathway in *S. Cerevisiae*. The results indicate that Nessie can learn the dynamics of biochemical reaction networks for a wide range of parameters, allowing us to investigate physical properties of a given system such as multimodality and parameter identifiability. Furthermore, in Section 4.3, we demonstrate how Nessie enables us to build upon [26] and perform efficient parameter inference from population snapshot data, a challenging problem for CME-based models. We conclude by discussing our results in Section 5.

## 2 Background

### 2.1 Chemical Master Equation

This section aims to provide a brief review of the CME and its use to model stochastic reaction networks in biology. We refer to [6, 29] for a readable and comprehensive treatment of the theory.

A biochemical reaction network consists of species *X_i_* (*i* = 1,…, *N*) and *R* reactions of the form

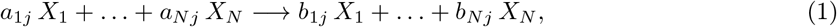

where *a_ij_* and *b_ij_* respectively denote the numbers of reactant and product molecules of species *i* in the chemical reaction *j*. The stoichiometric matrix is defined as *S_ij_* = *b_ij_* − *a_ij_* (the net change in the number of molecules of species *X_i_* when reaction *j* occurs). The state of the system is determined by the state vector ***n*** = (*n*_1_,…, *n_N_*), where *n_i_* is the number of molecules of species *X_i_* present in the system.

Starting with initial conditions *P*(***n**, t* = 0) =*P*_0_(***n***), the time evolution of the probability distribution over the states *P*(***n**, t*) is described by the CME:

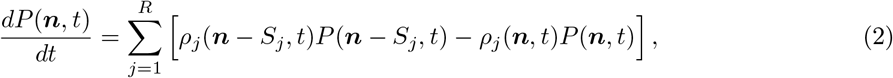

where *ρ_j_*(***n**, t*) is the propensity function of reaction *j* and *S_j_* is the *j*^th^ column of the stoichiometric matrix *S*. The propensity *ρ_j_*(***n**, t*) is the rate at which reaction *j* occurs when the system is in state ***n*** at time *t*; more formally, *ρ_j_*(***n**, t*)*dt* is the probability that reaction *j* will take place in the infinitesimally short time interval [*t, t* + *dt*) [30].

### 2.2 Neural Networks

A neural network learns a mapping *f* between inputs ***x*** and outputs ***y*** = *f*(***x***) via a parametric approximation *f_ϕ_* such that *f_ϕ_*(***x***) ≈ ***y***. The basic building block of a neural network is a single neuron, which performs the mapping ***x*** ↦ *g*(***x*** · ***w*** + *b*) for a weight vector ***w***, a bias *b* and a nonlinear activation function *g*. Several neurons arranged in parallel form a layer, and their outputs can be treated as inputs to another layer of neurons. By combining several layers in a row one gets a standard feedforward neural network, illustrated in Fig. 1. Here the first layer is called the input layer, the last layer is the output layer and the layers in between are hidden layers. For a comprehensive introduction to neural networks and deep learning we refer to [28].

**Figure 1:**
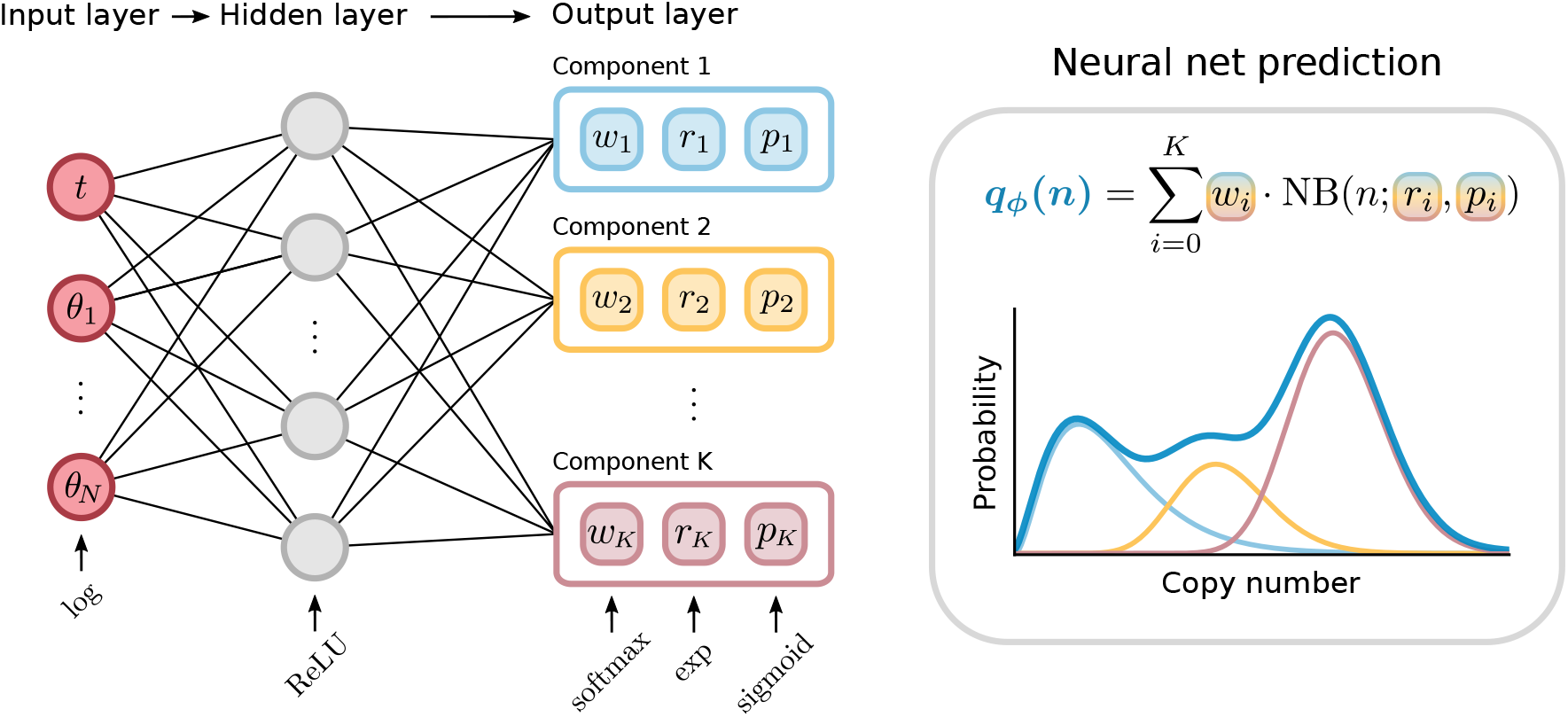
Architecture of Nessie, a simple feedforward neural network with one hidden layer. The input layer takes in the time *t* and the model parameters ***θ***, and the output layer returns the corresponding negative binomial mixture components. Activation functions are indicated at the bottom.

Using activation functions with each neuron enables neural networks to learn complicated nonlinear mappings. Commonly used activation functions in the Machine Learning community are sigmoid functions, Rectified Linear Units (ReLUs) [31] and variants thereof [28]. For these activation functions one can show that a feedforward neural network with a single hidden layer and a sufficient number of neurons acts as a universal approximator, i.e. it is able to represent any sufficiently smooth function [32] to arbitrary accuracy. In theory, a better approximation could be achieved using a deep neural network with multiple hidden layers, which can compose simpler functions into increasingly more complex ones. Such “deep” neural networks, often combining a variety of architectures more complex than a simple feedforward neural network, can outperform their shallow counterparts on difficult tasks such as Natural Language Processing, Computer Vision and others [28], which has led to a surge of interest in Deep Learning in recent years [13–16, 33]. As we will see, however, a single hidden layer is enough for our purposes.

The network parameters *ϕ* (weights and biases for each neuron) that minimize the discrepancy between the mapping *f_ϕ_* represented by the neural network and the true mapping *f*, are not known and have to be learned from data. This is commonly achieved by constructing a labeled set of training data 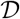 containing *N* ≫ 1 different input-output pairs, and minimizing a *loss function* 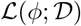 that measures the deviation between the neural mapping *f_ϕ_* and the target:

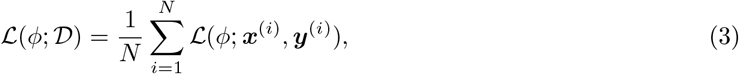

where (***x***^(*i*)^, ***y***^(*i*)^) denotes the *i*^th^ example pair in the training dataset. The most appropriate loss function depends on the type of data and the task the neural network is trying to perform: common examples are L2 distances for regression, cross-entropy for classification and negative log-likelihoods for inference problems [28].

As the loss function is often highly nonlinear and nonconvex, minimizing it with respect to the network parameters *ϕ* is a difficult task, most often performed using iterative gradient-based optimizers [34]. This requires the loss function to be differentiable with respect to the weights. Computing the gradients of the loss function is usually done using the backpropagation algorithm [28], which is implemented in most common deep learning frameworks such as Flux [35] or PyTorch [36]. Once the gradients have been computed one can use an optimization algorithm such as stochastic gradient descent or Adam [37] to minimize the loss.

In practice the behavior of a neural network and its training procedure are determined by a number of hyperparameters, such as the number and size of hidden layers, the activation functions applied on each layer, and the choice of the optimizer (as well as its associated parameters). There is no universal formula for determining the best hyperparameter choices for each task, and hence one has to resort to heuristics and hyperparameter tuning to find the best setup, which can be one of the most time-consuming aspects of training complicated neural networks. We discuss these practical considerations in connection to our approach in Appendix A.

Note that minimizing the loss function over the training data does not guarantee that the neural network will be able to *generalize*, i.e. accurately learn the mapping for previously unobserved inputs. For this reason, the network’s generalization ability is usually evaluated on a separate validation dataset made up of examples that are not included in the training data [28]. Comparison of the loss on the training and validation datasets during the training procedure allows us to perform effective hyperparameter tuning. Finally the predictive performance of the trained network can be accurately measured on a separate test dataset consisting of yet another set of input examples.

## 3 Nessie

Our goal in this paper is to learn marginal distributions predicted by the CME for different parameters and measurement times. As such the inputs to our neural network will consist of the chemical reaction network parameters ***θ*** and the time *t*; as these can span several orders of magnitude we log-transform them first. Although we work with fixed initial conditions for each reaction system, this constraint could be relaxed by adding the molecule numbers at time *t* = 0 as extra inputs to the neural network.

We approximate the marginal distribution of interest by a mixture of negative binomials, a flexible parametric class of distributions that has been shown to be very accurate for a large class of reaction networks [25, 26]. Indeed, it is known that single-time marginal distributions predicted by the CME for many different reaction networks can be modeled as a mixture of negative binomials in the presence of timescale separation [25, 38–41]. Experimental measurements of mRNA and protein distributions in bacterial, yeast and mammalian cells show that these are often fit well by such mixtures, even when timescale separation is not applicable [42–45]. We remark that a mixture of negative binomials always has a Fano factor (variance over mean) greater than 1, and systems whose Fano factor is significantly less than 1 (see e.g. [46]) would benefit from a different parametric approximation which we shall not consider here.

A mixture of negative binomials can be parameterized as

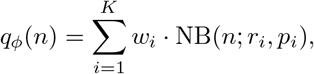

where *K* is the number of mixture components, *w_i_* is the weight of the *i*-th component and NB(*r_i_, p_i_*) is a negative binomial distribution with parameters (*r_i_, p_i_*). The number of components is fixed *a priori* and the weights are subject to the normalization constraint *w*_1_ + … + *w_K_* = 1. Our task is therefore to learn a mapping 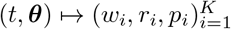 from the input parameters to those of the output distribution.

Each parameter characterizing the output distribution is represented by a single neuron in the output layer. To respect the constraints on the weights *w_i_* we apply the *softmax* activation function to the corresponding neurons: the outputs are exponentiated, then normalized to sum to 1. For the neurons corresponding to the count parameters *r_i_* we choose exponential activation functions, and for the probabilities *p_i_* we use sigmoid activation functions. The architecture of our neural network is shown in Fig. 1.

The number of hidden layers and the number of neurons per hidden layer can have a large impact on the representational power of a neural network, as noted in Section 2.2. The networks we build throughout this paper contain only a single hidden layer as we have found such architecture in our case to be easier to train and provide better predictive performance than “deeper” networks (see Section B for more details), an observation corroborated in [18]. We choose the number of neurons in the hidden layer depending on the complexity of the chemical reaction network at hand and use the ReLU activation function as it enables efficient training [31].

In our setup a single training point consists of an input point ***x*** = (*t, θ*) and a reference distribution *p* of target molecules at time *t* for the specified reaction network with parameters ***θ***, obtained by averaging over a number of SSA trajectories (or by using the FSP). We build the training set by sampling parameter sets ***θ*** in the parameter region of interest and running simulations at each ***θ***; this can be done in parallel for all parameters. In order to ensure that the training data evenly cover the entire parameter region we use Sobol sequences [47], which generally provide more uniform coverage than random sampling.

A common method to match distributions in the statistics literature is to minimize the Kullback-Leibler (KL) divergence *D*_KL_(*p* ∥ *q_ϕ_*), where *p* is the target distribution and *q_ϕ_* the prediction; this procedure is mathematically equivalent to maximizing the average log-likelihood under *q_ϕ_* of a sample drawn from *p* [28]. Hence we use the KL divergence as our loss function (cf. Eq. 2.2) for each point in the training set:

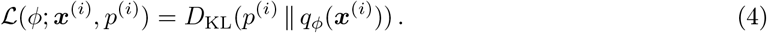

This is equivalent to the cross-entropy, up to the addition of a constant that does not depend on the network weights *ϕ*. Computing the mixture of negative binomials *q_ϕ_* for a given input point is straightforward using Eq. 4. The complete workflow for training Nessie is shown in Figure 2.

**Figure 2:**
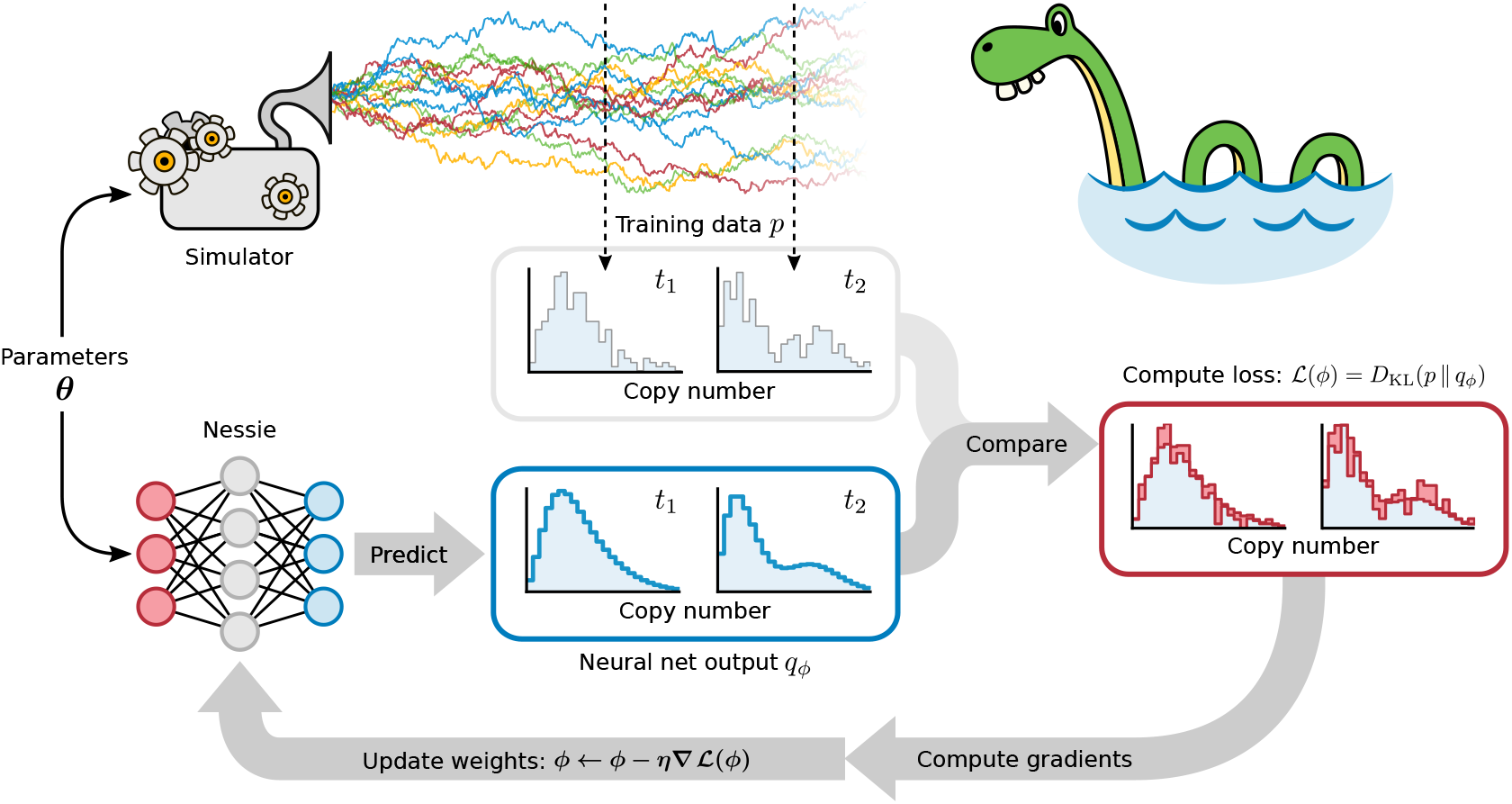
Workflow for training Nessie. Given model parameters ***θ***, the reaction network is simulated repeatedly using the SSA to obtain empirical distributions at training time points *t*_1_, *t*_2_,…. These are then compared with the output of the neural network and the total loss 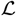 is computed. In order to decrease the loss, the neural weights *ϕ* are updated iteratively via gradient descent until the loss has converged. In the figure *η* denotes the learning rate (see Appendix A) and *D*_KL_(*p* ∥ *q_ϕ_*) the KL divergence between the true distribution *p* and the neural network prediction *q_ϕ_*.

We note that maximizing the average log-likelihood for a mixture of Gaussians, as is commonly done with Mixture Density Networks (MDNs) [24], can lead to stability issues for more than one component. For example, the neural network can learn to place a Gaussian component at zero with arbitrarily low variance, which will give an arbitrarily high likelihood if 0 occurs anywhere in the training dataset, irrespectively of the overall quality of fit — an example of overfitting (see Appendix A). This is because in the continuous case one deals with probability *densities*, which can become arbitrarily large in contrast to probabilities. We believe that this phenomenon is responsible for some common numerical issues observed e.g. in [21, 27, 28]. One can attempt to remedy this issue by integrating the density over a finite interval (say, [−0.5, 0.5]), or by regularizing the precision of each component (thereby adding hyperparameters to the training procedure). In contrast, mixtures of negative binomials were not prone to overfitting in our experiments and did not require any form of regularization.

In what follows, we quantify the relative accuracy of the trained neural networks by computing the Hellinger distance between the predicted and test distributions. Although the KL divergence is more suited as a loss function [28] to train the neural network for its computational efficiency, the Hellinger distance is a bounded metric and a more interpretable measure of the model’s predictive performance.

We used Julia with `Flux.jl` [35] to implement neural networks. Gradients of the loss function were computed directly by Flux using the built-in `Zygote.jl` automatic differentiation system [48]. The training datasets were constructed by defining chemical reaction networks via `Catalyst.jl` and simulating them using `DifferentialEquations.jl` [49] (SSA) and `FiniteStateProjection.jl` (FSP). All numerical experiments were performed on a Intel Xeon Silver 4114 CPU (2.2 GHz) using 16 threads.

## 4 Results

### 4.1 Autoregulatory Feedback Loop

We first consider a simple autoregulatory feedback loop illustrated schematically in Fig. 3**a**. This system contains a single gene with two promoter states *G_u_* and *G_b_*, each associated with different protein *P* production rates *ρ_u_* and *ρ_b_* (mRNA dynamics are not modeled explicitly). The feedback is introduced via reversible binding of a protein molecule to the promoter region with binding rate *σ_b_* and unbinding rate *σ_u_*, which causes switching between the two promoter states. Finally, protein degradation is modeled by an effective first-order reaction. This system is a rudimentary example of stochastic self-regulation in a gene: the model functions as a positive feedback loop if *ρ_b_* > *ρ_u_*, and a negative feedback loop if *ρ_b_* < *ρ_u_*.

**Figure 3:**
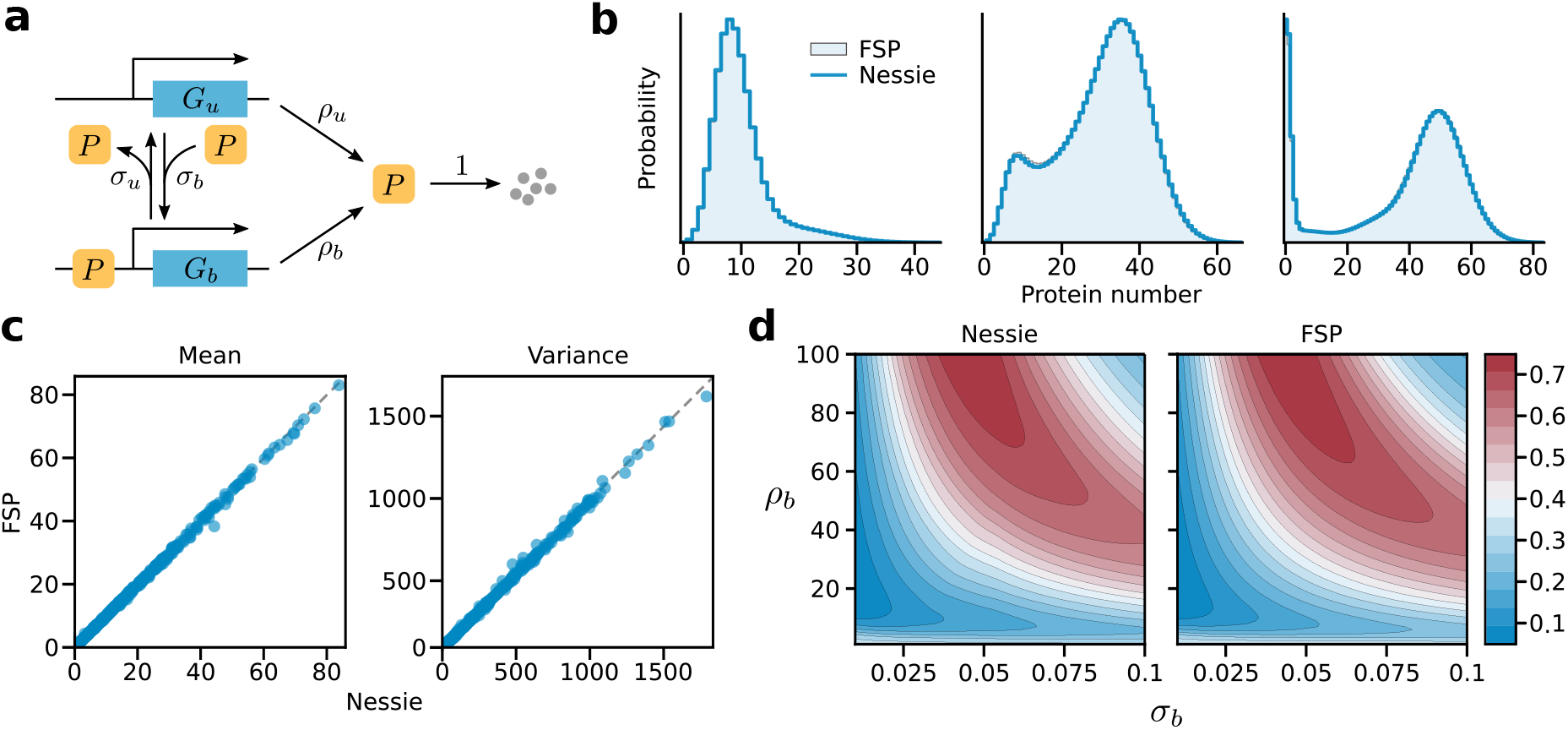
Nessie applied to an autoregulatory genetic feedback loop. **a)** Schematic of the reaction network. We assume mass action kinetics for all reactions. **b)** Protein distributions for three different test parameter values (indicated in Table 1). The ground truth distributions were computed using the FSP. **c)** Comparison of true and predicted means and variances of protein numbers at time *t* = 100 for the test set containing 500 parameter values. True means and variances were again computed using the FSP. **d)** Exact and predicted bimodality coefficients as a function of *ρ_b_* and *σ_b_*, where we set *σ_u_* = *ρ_u_* = 1 and *t* = 100. Here the bimodality coefficients predicted by Nessie closely agree with their ground truth values.

Although the CME of the autoregulatory feedback loop has only been solved analytically in the steadystate [50], an efficient time-dependent numerical solution can be obtained with the FSP in this specific case as the model contains few molecular species and chemical reactions. Estimating the probability distributions for the autoregulatory feedback loop via the FSP is much faster than using the SSA. This makes the autoregulatory feedback loop an ideal toy model for our initial experiments as we can relatively quickly build arbitrarily large training datasets with the FSP, calibrate the neural network, and probe the performance of Nessie in capturing the marginal distributions of protein numbers.

We use a training set of size 1k, a validation set of size 100 and a test set of size 500, sampled using a Sobol sequence in the parameter region indicated in Table 1. For each datapoint we take four snapshots at times *t* = {5, 10, 25, 100} and construct the corresponding histograms using the FSP. Our neural network consists of a single hidden layer with 128 neurons and outputs 4 negative binomial mixture components; we use a batch size of 64 for training. More details on the training procedure and the hyperparameter choices are given in the Appendices A and B.

**Table 1:** Parameters and parameter ranges used for the autoregulatory feedback loop presented in Section 4.1. The initial conditions are zero proteins and the gene in the unbound state *G_u_*.

| Parameter | $t$ | $\sigma_u$ | $\sigma_b$ | $\rho_u$ | $\rho_b$ |
| --- | --- | --- | --- | --- | --- |
| Range | - | 0-2 | 0-0.1 | 0-10 | 0-100 |
| Fig. 3 b)-(i) | 10 | 0.94 | 0.01 | 8.40 | 28.1 |
| Fig. 3 b)-(ii) | 25 | 0.69 | 0.07 | 7.20 | 40.6 |
| Fig. 3 b)-(iii) | 100 | 0.44 | 0.08 | 0.94 | 53.1 |
| Fig. 4 b) | 10 | 1.17 | 0.05 | 6.90 | 47.9 |

In Fig. 3**b** we show the protein distributions for three test parameters sets, comparing Nessie predictions to the FSP results. Our approach provides highly accurate fits for the different distribution shapes obtained at these points in the parameter space, showcasing the flexibility of negative binomial mixtures in approximating the CME of the autoregulatory feedback loop.

Having learned the output distributions for this chemical system we can compute various quantities of interest from these. In Fig. 3**c**, we compare the means and variances of the protein number predicted by Nessie to the true values computed using the FSP for all points in the test dataset. Furthermore, in Fig. 3**d** we analyse the bimodality coefficient as defined in [51]. The bimodality coefficient of a distribution is defined as 1/(*κ* − *γ*^2^), where *κ* and *γ* are the skewness and kurtosis respectively, and is a measure of bimodality with higher values corresponding to strongly bimodal distributions. We see that Nessie provides a good approximation to this quantity and closely matches the FSP results.

Numerically estimating the bimodality coefficient for many different parameters is a computationally intensive task, whereas predicting it using the neural network, once trained on its 1k datapoints, is very quick: using Nessie we can produce the plotted heatmap in 0.03 s, in contrast to the FSP which takes 240s. This example illustrates how we can apply the neural network to rapidly and efficiently analyze large swathes of parameter space and, in turn, to determine the regions of bimodality in the system. Similarly, our approach scales to more complicated chemical reaction networks that are not amenable to study using the FSP, allowing us for example to perform global sensitivity analysis for the genetic toggle switch in Section 4.2.

Finally, in Fig. 4 we consider using the SSA as an alternative to the FSP for constructing the training dataset. As each training histogram is built from a number of SSA samples, we investigate how many samples per training point are required to accurately train the network. Note that the numerical FSP solution is virtually indistinguishable from the exact CME solution as its approximation error can be systematically reduced by increasing the size of the truncated state space [7], and hence it is effectively equivalent to an infinite number of SSA trajectories. We see in Fig. 4**a** that for the autoregulatory feedback loop using as few as 100 trajectories per training point enables Nessie to produce good approximations of the true distributions. In contrast, obtaining similar quality fits using the SSA alone would require two orders of magnitude more samples per parameter, as demonstrated in Fig. 4**b** where we compare the histograms obtained using 100, 1k and 10k SSA trajectories. One may expect this effect to be amplified for larger system sizes and more widely spread out distributions, where many more simulations are typically needed to obtain smooth histograms.

**Figure 4:**
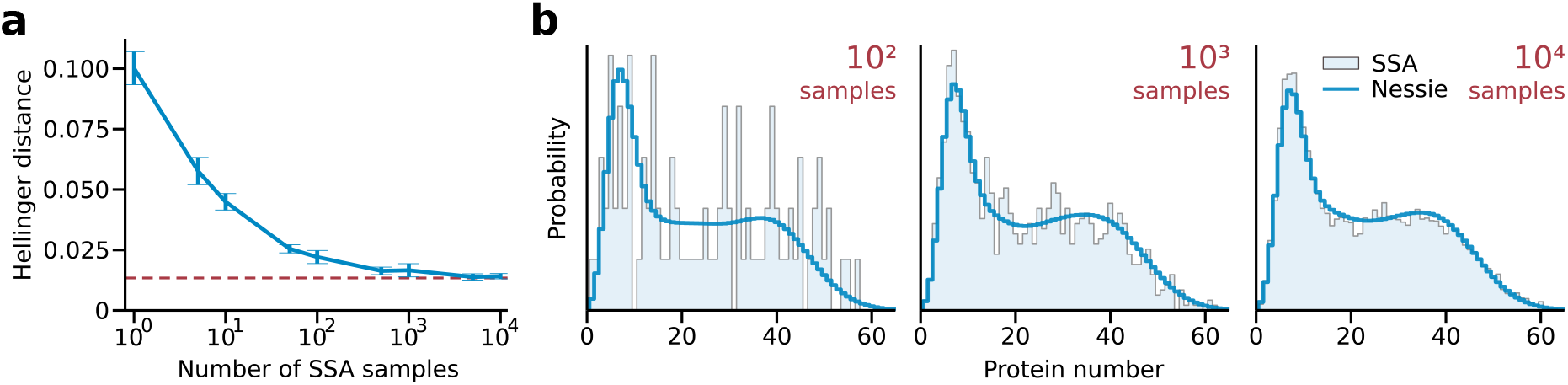
Training Nessie using the SSA. **a)** Mean Hellinger distance computed over the validation dataset versus the number of SSA trajectories used to construct the histogram of each training datapoint. Here the validation dataset consists of 100 different parameter values at 4 time snapshots, constructed with the corresponding number of SSA trajectories. The error bars are obtained by averaging over 10 independent Nessie training runs, where for each run we resample the training and validation datasets and train a new model, as done for the manual tuning of other hyperparameters discussed in Appendix B and Fig. 8. The red dashed line indicates the accuracy of Nessie trained on the FSP data. **b)** Example distributions constructed with 100, 1k and 10k SSA samples (indicated in red at the top) compared to Nessie predictions. Note that Nessie is retrained for each plot using the respective number of trajectories. The parameter values of the feedback loop are given in Table 1.

### 4.2 Genetic Toggle Switch

For our next experiment we consider a stochastic model of the genetic toggle switch, one of the first synthetic biological circuits [52]. The reaction network, introduced in [53], is composed of two mutually interacting genes (which we label *A* and *B*) and takes into account transcription, translation and the subsequent degradation of the produced mRNA and proteins (see Fig. 5**a**). The translated protein A can bind to the promoter region of the gene that produces protein *B*, and vice versa with protein *B* binding to the gene promoter of protein *A*. This results in effective transcriptional regulation: depending on the mRNA production rate associated with each promoter state, the process can either lead to repression or activation of transcription for each species. In addition, the system contains post-transcriptional regulation mediated by protein A binding to the mRNA of species B and modulating its translation accordingly [54].

**Figure 5:**
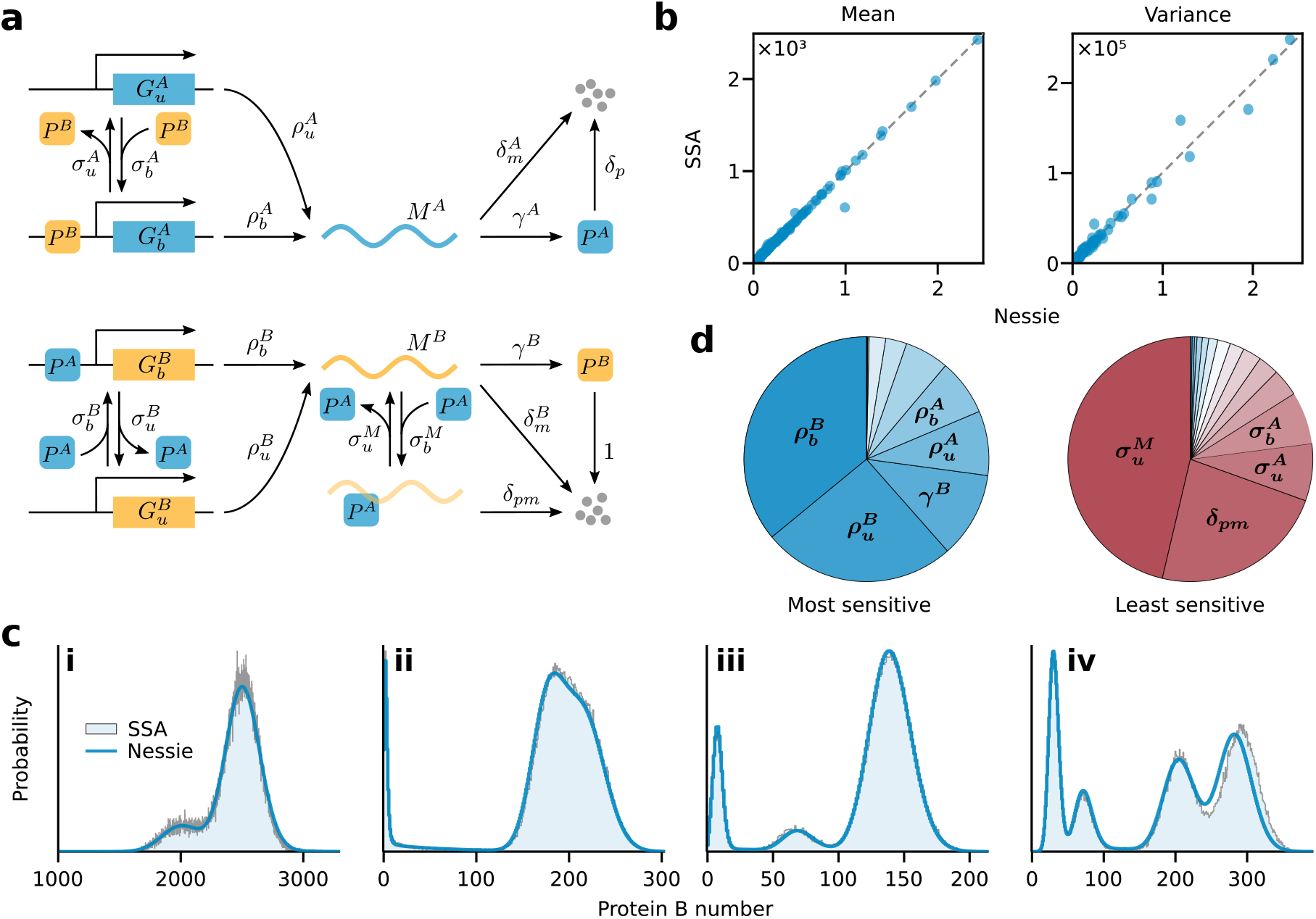
Nessie applied to a genetic toggle switch with post-transcriptional regulation. **a)** Schematic of the reaction network (assuming mass action kinetics for all reactions). **b)** Comparison of true and predicted means and variances of the protein *B* numbers for the test set consisting of 500 different parameter values at time *t* = 100 constructed using 100k SSA trajectories. **c)** Protein B distributions for four different test parameter values (specified in Table 2). The SSA distributions were computed by averaging over 100k trajectories. **d)** Sensitivity of the Fano factor of protein *B* to parameter perturbation at time *t* = 100, where the pie charts show the most and least sensitive parameters. The results are obtained by using Nessie to compute the logarithmic sensitivity of the Fano factor to the 16 model parameters for 100k parameter values drawn from a Sobol sequence covering the training range given in Table 2. We observe that only a few reaction parameters can be identified as typically the most/least sensitive (indicated in bold).

The toggle switch is noticeably more intricate than the autoregulatory feedback loop considered in Section 4.1. It exhibits rich dynamics highlighted by diverse protein distributions that can be highly multimodal [53]. Due to the considerable number of reaction parameters, the frequent occurrence of high copy numbers (> 1000) and the complexity of the observed distributions, studying this system poses significant problems both analytically and computationally. This makes it a good challenge for Nessie.

Our aim is to predict the probability distributions of the target protein B in the genetic toggle switch. Note that we could similarly consider the distribution of protein A or, alternatively, the neural network architecture itself could be extended to predict both marginal distributions for the two proteins in parallel. If however joint distributions are required, the univariate mixture of negative binomials we use needs to be replaced by a multivariate equivalent, a topic which we do not pursue in this paper.

We draw 40k, 100 and 500 parameters for the training, validation and test datasets respectively using a Sobol sequence in the parameter region indicated in Table 2. For each parameter set we take 8 snapshots at times *t* = {2, 4, 10, 16, 32, 50, 74, 100}. The complexity of the system prevents us from using the FSP to construct the reference histograms and hence we resort to the SSA. As discussed in Section 4.1, a relatively small number of SSA samples can be used to successfully train the neural network. For this reason, we use 1k simulations for each training datapoint and 100k simulations for the validation and test data (in order to ensure a more accurate comparison to the true distributions). In this case, we use a neural network with a single hidden layer of 1024 neurons and 6 output mixture components, and fix the batch size to 1000 for the training procedure (see Appendix A for more details). The remainder of our setup is the same as in the previous example.

**Table 2:** Parameters and parameter ranges for the genetic toggle switch presented in Section 4.2. The initial conditions are zero proteins/mRNAs and both genes in the unbound states 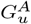 resp. 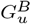.

| Parameter | $t$ | $\sigma_b^B$ | $\sigma_u^B$ | $\sigma_b^A$ | $\sigma_u^A$ | $\rho_u^A$ | $\rho_b^A$ | $\gamma^A$ | $\delta_m^A$ | $\delta_p$ | $\rho_u^B$ | $\rho_b^B$ | $\gamma^B$ | $\delta_m^B$ | $\sigma_u^M$ | $\sigma_b^M$ | $\delta_{pm}$ |
| --- | --- | --- | --- | --- | --- | --- | --- | --- | --- | --- | --- | --- | --- | --- | --- | --- | --- |
| Range | - | $0-5 \times 10^{-4}$ | 0-0.1 | $0-5 \times 10^{-4}$ | 0-0.1 | 0-500 | 0-500 | 0-12 | 1-20 | 0-2 | 0-500 | 0-500 | 0-12 | 1-20 | 0-2 | 0-0.5 | 0-100 |
| Fig. 5 b)-(i) | 35 | $5 \times 10^{-4}$ | 0.070 | $1.6 \times 10^{-4}$ | 0.0680 | 367 | 119.9 | 5.3 | 14.36 | 1.35 | 413 | 472.3 | 11.9 | 1.74 | 0.98 | 0.43 | 65.6 |
| Fig. 5 b)-(ii) | 16 | $2 \times 10^{-4}$ | 0.060 | $4.8 \times 10^{-4}$ | 0.0095 | 328 | 190.2 | 11.1 | 15.84 | 1.01 | 467 | 11.3 | 8.7 | 13.91 | 1.14 | 0.17 | 92.2 |
| Fig. 5 b)-(iii) | 70 | $3 \times 10^{-4}$ | 0.018 | $0.2 \times 10^{-4}$ | 0.0200 | 222 | 474.9 | 4.0 | 6.96 | 0.68 | 360 | 82.1 | 4.2 | 7.13 | 0.10 | 0.41 | 68.4 |
| Fig. 5 b)-(iv) | 100 | $2 \times 10^{-4}$ | 0.005 | $3.5 \times 10^{-4}$ | 0.0460 | 371 | 69.1 | 7.1 | 13.90 | 0.87 | 128 | 452.7 | 8.4 | 11.98 | 1.10 | 0.20 | 72.6 |
| Fig. 6 | - | $4 \times 10^{-4}$ | 0.060 | $2.2 \times 10^{-4}$ | 0.0034 | 270 | 61.3 | 8.0 | 9.46 | 1.78 | 308 | 351.2 | 10.1 | 12.27 | 0.09 | 0.38 | 85.6 |

In Fig. 5**b** we verify that the moments (means and variances) of the protein *B* numbers predicted by Nessie closely match those computed using the SSA for all test datapoints. Furthermore, in Fig. 5**c**, we compare the predicted protein distributions to the true distributions constructed by averaging over 100k SSA realizations. Notably, the trained neural network is able to reconstruct the complex distributions and provides a good approximation to CME solution of the genetic toggle switch. The promising performance of Nessie highlights the usefulness of neural emulators for dealing with stochastic chemical reaction networks that go beyond the more tractable examples that are typically studied in the literature.

Next we explore the sensitivity of the genetic toggle switch to noise. The Fano factor, which is defined as the ratio of the variance of molecule numbers to the mean molecule number, is a commonly used measure of deviations from Poisson noise and the extent of transcriptional/translational bursting. We investigate how the Fano factor of the protein B number changes upon parameter perturbation. Using the trained neural network we have computed the logarithmic sensitivity [55] of the Fano factor of protein B to all reaction parameters over a wide parameter range and identified the most and least sensitive parameters on average, as shown in the pie charts in Fig. 5**d**. In particular, we can identify a few parameters that are the most or least sensitive to noise in the majority of cases. For example, for over 60% of the parameter space, the mRNA production rates 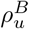 and 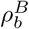 are the most sensitive, and hence tweaking these parameters is usually the optimal way to control the fluctuations in the protein B number. Note that performing such global sensitivity analysis is highly computationally expensive using the SSA, whereas with Nessie it can be approximated within minutes, making it possible to significantly accelerate further parameter exploration studies.

Although we trained Nessie with distribution snapshots only at a few fixed time points, an obvious question of interest is whether the neural network can capture the temporal dynamics of the chemical system over its whole trajectory. In Fig. 6 we plot the Hellinger distance between the predicted and true (SSA) distributions at times *t* = {1, 2,…, 100} averaged over the 500 test parameter sets. Remarkably, the predictive accuracy is largely similar throughout the whole time range, indicating that the neural network is able to effectively interpolate between the time points it has seen during training. Note that the worse performance at *t* < 2 is expected as we do not train on the initial transient during which the dynamics rapidly evolve from the initial condition.

**Figure 6:**
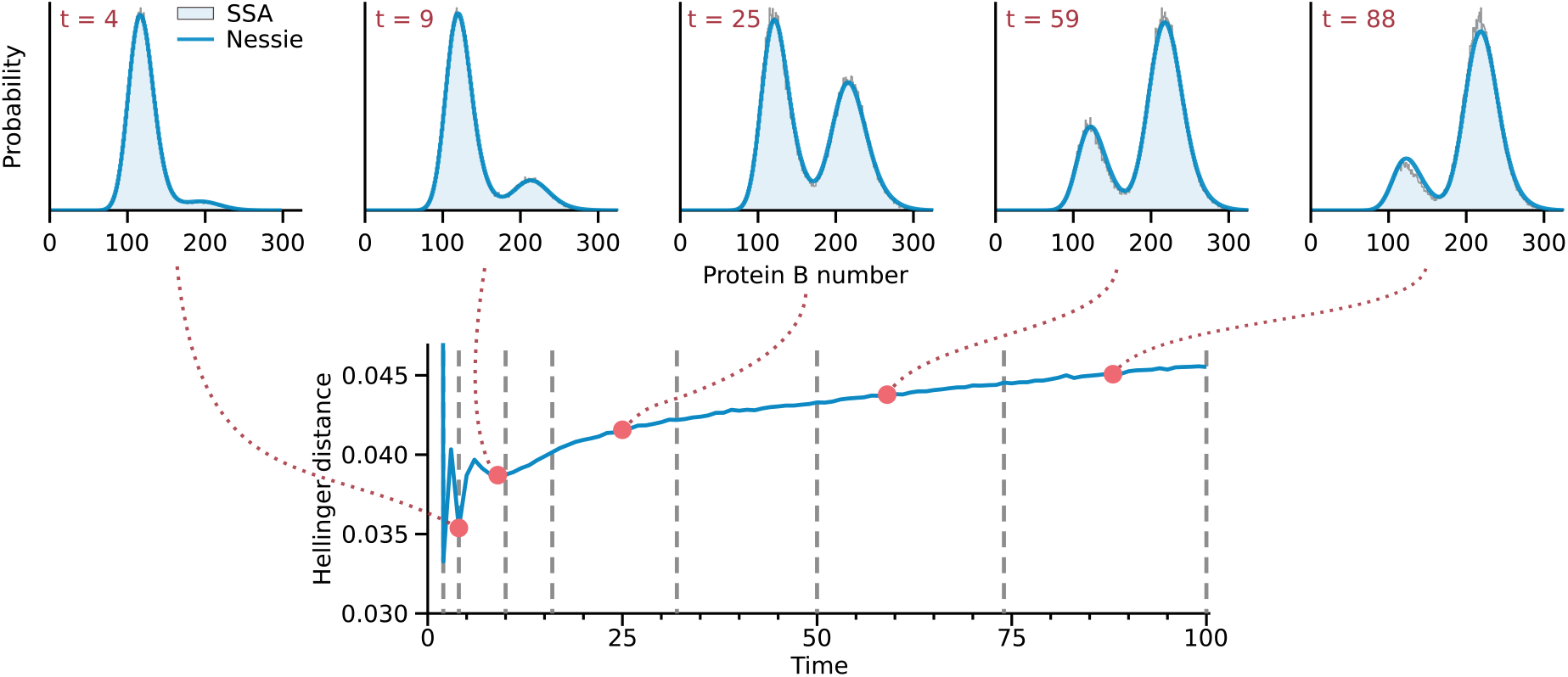
Using Nessie to interpolate and predict the protein *B* dynamics in time. Bottom: Mean Hellinger distance computed over the test dataset made up of 500 different parameter values (constructed using 100k SSA trajectories) evaluated at times *t* = {1, 2,…, 100}. The vertical gray dashed lines indicate the time snapshots used for training the neural network, showing that the predictive error does not notably increase in between the training points. Top: Time evolution of the protein distribution predicted by Nessie and the SSA (averaged over 100k trajectories) for an example parameter set given in Table 2, demonstrating Nessie’s ability to accurately capture the time evolution of the protein *B* distribution.

### 4.3 MAPK Pathway

We finally apply Nessie to a biological model of the MAPK pathway in *S. Cerevisiae* with the aim of inferring system parameters using experimental data from [56]. The reaction network can be seen in Fig. 7**a** and is modified from [56], removing extrinsic noise contributions. It describes the pSTL1 gene and includes activation due to a time-varying *hog1* signal, chromatin remodeling, transcription and translation. When yeast is subjected to external osmotic pressure, activation of the MAPK signaling cascade results in doubly phosphorylated *hog1* molecules entering the nucleus. These bind to the pSTL1 promoter, which is initially in an inactive state (*G*\*\*). Upon binding of *hog1* to the gene, subject to chromatin structure remodeling via the chromatin remodeling complex (*CR*), starts transcribing mRNA, which after translocation into the cytosol is translated into protein. In [56] the authors used flow cytometry to measure protein number distributions for the MAPK pathway, for which they proposed the above model. Protein distributions predicted by this model tend to be bimodal with a sharp peak at 0, as depending on the parameters a sizeable fraction of cells never starts transcribing mRNA before the *hog1* signal decays.

**Figure 7:**
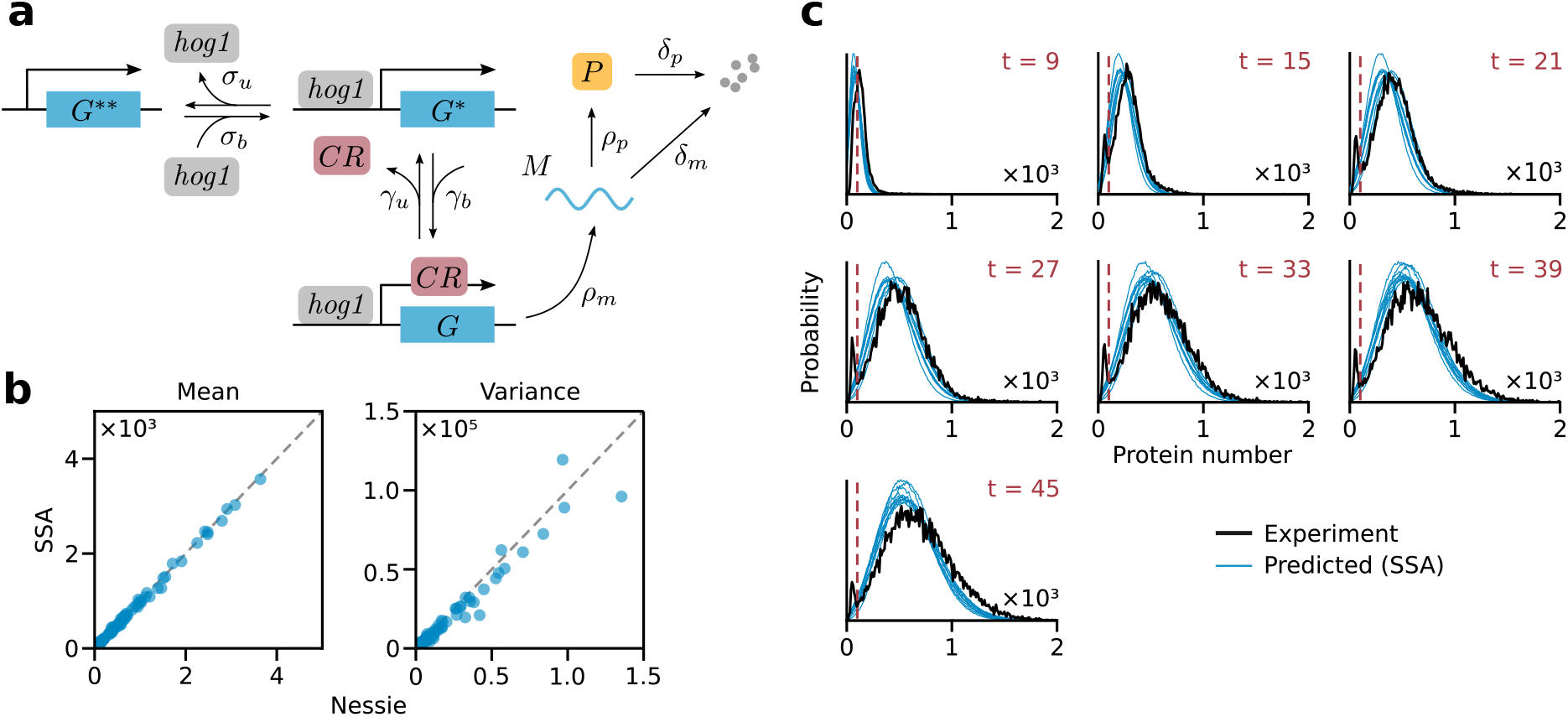
Results for the MAPK pathway example. **a)** Schematic of the reaction network (see Appendix C.3 for details). The *hog1* concentration over time is taken from [56]. **b)** True and predicted means and variances of the protein distribution for the test set consisting of 100 different parameter values at time *t* = 27 constructed using 100k SSA trajectories. **c)** Comparison of the experimental distribution (black) with those predicted by the CME for the parameters inferred using Nessie (blue). To probe parameter uncertainty we performed 10 independent estimation rounds; the resulting parameter values are given in Table 2). SSA distributions were computed by averaging over 1000k trajectories.

The data measured in [56] consists of intensity measurements (in arbitrary units, AU) at times *t* = {3, 9, 15, 21, 27, 33, 39, 45} (in minutes) after salt was added to the solution to induce osmotic shock in the cells, which triggers the MAPK pathway. Since the fluorescence intensity per protein (*I/P*) was not measured in these experiments we assumed a value of 1AU per protein and rounded the estimated protein number to the nearest integer (in [56] it was noted that identifying all parameters is not possible from these experiments).

Observing that the experimental distributions at most times had a peak near 0 whose width was consistent across time points, we binned all observations less than 100AU, the approximate width of the peaks. The observed peaks are best explained by measurement noise that does not allow us to identify the exact protein numbers in the low copy number regime. Binning in this case, while potentially losing some information, renders the procedure more reliable than the alternative of discarding observations below the threshold [57, 58]. Since measurements at time *t* = 3 were almost entirely below the threshold we discarded that time point for inference purposes.

In order to fit parameters we minimized the total Hellinger distance between the experimentally observed distribution and the model output at each time point as predicted by Nessie, treating all observations below 100 as lying in one bin. The training, validation and test sets consist of 15k, 250 and 100 points, respectively, which were generated from a logarithmic Sobol sequence in the parameter region indicated in Table 3, chosen around the maximum a posteriori estimates reported in [56]. We use 1k simulations for each training datapoint and 100k simulations for the validation and test data. The size of the hidden layer was set to 2048 and the number of mixture components was 5. Since a significant fraction of the simulated trajectories had 0 proteins we added a sixth component that was set to be a Dirac delta at 0; this was performed by adding a single output neuron predicting the weight of this peak. To speed up training we split the procedure into two rounds, first training with 10% of the training data for 100 rounds and then with the entire training set for 400. We used a batch size of 1k throughout. Fig. 7**b** shows that the means and variances predicted by the resulting neural network are close to those obtained using the SSA.

**Table 3:** Search ranges and 10 estimated parameter sets for the MAPK pathway presented in Section 4.3. The initial conditions were zero mRNA and proteins, 225 *CR* molecules and the gene in the deactivated state *G*\*\*.

| Parameter | $b$ | $h$ | $K_m$ | $V_{\max}$ | $\eta_u$ | $\sigma_b$ | $\sigma_u$ | $\rho_m$ | $\rho_p$ | $\delta_m$ | $\delta_p$ |
| --- | --- | --- | --- | --- | --- | --- | --- | --- | --- | --- | --- |
| Range | 0.001–0.1 | 1–10 | 0.01–1 | 0.1–10 | 0.1–10 | $10^{-4}$–$10^{-2}$ | 0.001–0.1 | 0.1–1 | 0.001–0.01 | $10^{-4}$–$10^{-3}$ | $10^{-5}$–$10^{-3}$ |
| Fig. 7 b) | 0.0070 | 7.3 | 0.196 | 3.16 | 3.12 | $28.19 \times 10^{-4}$ | 0.054 | 0.555 | 0.00661 | $4.31 \times 10^{-4}$ | $45.4 \times 10^{-4}$ |
| | 0.0186 | 3.7 | 0.184 | 0.95 | 2.92 | $8.17 \times 10^{-4}$ | 0.027 | 0.353 | 0.00906 | $7.68 \times 10^{-4}$ | $28.7 \times 10^{-4}$ |
| | 0.0101 | 1.5 | 0.078 | 0.71 | 2.09 | $2.29 \times 10^{-4}$ | 0.008 | 0.456 | 0.00518 | $9.73 \times 10^{-4}$ | $4.97 \times 10^{-4}$ |
| | 0.0182 | 3.1 | 0.227 | 1.16 | 1.21 | $5.87 \times 10^{-4}$ | 0.036 | 0.334 | 0.00982 | $3.88 \times 10^{-4}$ | $2.01 \times 10^{-4}$ |
| | 0.0315 | 8.7 | 0.190 | 1.63 | 0.31 | $3.81 \times 10^{-4}$ | 0.025 | 0.368 | 0.00757 | $8.48 \times 10^{-4}$ | $45.3 \times 10^{-4}$ |
| | 0.0056 | 1.5 | 0.072 | 0.86 | 2.20 | $2.18 \times 10^{-4}$ | 0.010 | 0.394 | 0.00605 | $6.73 \times 10^{-4}$ | $5.22 \times 10^{-4}$ |
| | 0.0177 | 7.5 | 0.175 | 0.41 | 2.24 | $56.15 \times 10^{-4}$ | 0.047 | 0.507 | 0.00634 | $2.62 \times 10^{-4}$ | $2.19 \times 10^{-4}$ |
| | 0.0252 | 5.8 | 0.232 | 0.68 | 0.27 | $8.06 \times 10^{-4}$ | 0.031 | 0.459 | 0.00632 | $2.66 \times 10^{-4}$ | $4.70 \times 10^{-4}$ |
| | 0.0108 | 1.9 | 0.075 | 0.55 | 1.47 | $2.06 \times 10^{-4}$ | 0.008 | 0.381 | 0.00584 | $7.54 \times 10^{-4}$ | $5.22 \times 10^{-4}$ |
| | 0.0278 | 3.6 | 0.300 | 3.56 | 3.40 | $12.88 \times 10^{-4}$ | 0.055 | 0.453 | 0.00982 | $4.09 \times 10^{-4}$ | $1.45 \times 10^{-4}$ |

To evaluate the accuracy of the estimation scheme we ran the SSA at the predicted parameters, verifying that the results match the experimentally observed distribution (see Fig. 7**c** for the results using 10 estimated parameter sets). As can be seen in the figure our results do not reproduce the peak near 0 found in the experimental input, and an extensive parameter search did not lead to any parameters which exactly reproduce the experimental data. We therefore suspect that the model we used does not fully describe the dynamics of *hog1*-mediated gene expression and that obtaining better results will require a more detailed model than the one we are using.

While neural networks could be used to perform maximum likelihood estimation or Bayesian inference as in [59], we did not pursue likelihood-based approaches in this paper. Due to the large number of datapoints (over 100k), any small approximation error in the distributions predicted by Nessie will get amplified by several orders of magnitude: as the likelihoods for each datapoint add up to form the total likelihood of the data, the errors in the likelihood will, too. This leads to highly fluctuating likelihood values for similar parameters that are an artifact of the neural approximation and not present in the true model. This results in the predicted posterior being concentrated tightly around one parameter set where these fluctuations result in a marginally higher likelihood for the experimental data than the others, and the resulting uncertainty estimates reflect the approximation error incurred by Nessie instead of true parameter uncertainty. Such concentration of the estimated posterior due to randomness is a common problem in using MCMC with “tall data” [60], where estimating likelihoods for large datasets becomes very difficult.

Since fitting parameters using Hellinger distances does not directly provide uncertainty information, we estimated uncertainty by repeatedly fitting parameters to the experimental data; Table 3 shows the results of 10 fits. As can be seen in Fig. 7**b**), these results produce similar distributions under the CME, yet some parameters such as *σ_b_* and *δ_p_* are spread over an order of magnitude. Such parameter unidentifiability is common with the type of experimental data measured in biological experiments and should be taken into account when interpreting results. In particular the Hill coefficient, which was allowed to range from 1 to 10, could not be narrowed down within this range.

Once the network is trained, globally optimizing the Hellinger distance within the targeted parameter region takes a few minutes. Our approach should therefore be particularly suited for scenarios where distribution data is available for many copies of one network, e.g. when using a single gene expression model to analyse many different genes in an organism.

## 5 Discussion

In this paper we presented Nessie, a framework that allows us to train neural networks on simulation data to accurately estimate the solution of the CME for various biological systems. Our approach is scalable to complex nonlinear reaction networks with over a dozen parameters that exhibit diverse, multimodal dynamics across parameter space. We illustrated the performance of our approach on three examples: a well-studied autoregulatory feedback loop, the MAPK pathway in *S. Cerevisiae* and a complex genetic toggle switch. The latter two pose significant challenges both analytically and computationally due to the number of species and reactions involved, yet Nessie allows us to efficiently emulate them and analyze their properties, with applications for parameter exploration and estimation.

Nessie can be particularly useful in rapidly exploring large swathes of parameter space, for example to perform a local or global sensitivity analysis. This has many uses, e.g. to guide the tuning of parameters to find a desired phenotype [61], for the design of optimal experiments [62], to provide insights into the robustness and fragility tradeoff in genetic regulatory mechanisms [63]) and to find those parameters which most influence the size of transcriptional noise [64]. We note that performing any such analysis using the standard stochastic simulation methods like the SSA can be prohibitively computationally expensive [65].

Following methodology similar to that proposed in [59], Nessie can be used to fit models to data by matching experimentally observed distributions to those predicted by the neural network, as demonstrated in the case of the MAPK pathway model where we recovered model parameters that are mostly consistent with experimental observations. We remark, however, that this approach has to be used with care in the context of likelihood-based inference due to small approximation errors in the likelihood being amplified in the presence of many datapoints. In order to be reliable any such approach, including Bayesian inference, must take into account the bias introduced by the choice of approximation. This could be done e.g. by placing a prior over network weights and treating them as unobserved variables, sampling from the resulting Bayesian neural network using Hamiltonian Monte Carlo methods.

As discussed in Section 1, our approach differs from other studies that use neural networks to predict the dynamics of stochastic biochemical systems [19–23]. As all of these approaches try to learn different things, comparing them directly is not straightforward and is further complicated by the sensitivity of neural networks to architecture and hyperparameter choices [28]. An advantage of Nessie as presented in this paper is that it requires relatively little setup in terms of hyperparameter optimization. Due to its architectural simplicity a very limited amount of tuning is required, without requiring automated neural architecture search techniques [21], and we provide a detailed discussion of the relevant hyperparameter and training considerations in Appendices A and B. We hope that this will enable the interested reader to quickly deploy and apply Nessie to their favorite reaction network.

Although Nessie can relatively accurately interpolate in time between the training snapshots for such models like the genetic toggle switch, its performance may be inadequate when applied on systems with complex oscillatory behavior. This is a general limitation of our approach, which uses a simple feedforward network and therefore may not be able to efficiently represent oscillating functions. To remedy this, besides the recently proposed generative adversarial network-based approach in [23] one could consider more sophisticated neural network architectures such as recurrent neural networks [28] and universal differential equations [49]. This would allow us to extract temporal features such as power spectra and first passage times, which, while difficult to measure experimentally, have been shown to provide a wealth of information about the system and significantly aid in model discrimination [66–68].

## Data Availability Statement

Code for this paper is available at https://github.com/augustinas1/Nessie.

## Acknowledgments

This work was supported by the Alan Turing Institute Doctoral Studentship (under the EPSRC grant EP/N510129/1) for A. S., the EPSRC Centre for Doctoral Training in Data Science (EPSRC grant EP/L016427/1) and the University of Edinburgh for K. Ö. and a Leverhulme Trust grant (Grant No. RPG-2020-327) for R. G. The authors would like to thank Christoph Zechner for the courtesy of sharing his data for the MAPK pathway model.

## Appendix

### A Training Neural Networks

We train our neural networks using the Adam optimizer [37], one of the most popular optimization algorithms for this purpose. The gradients of the loss function with respect to the network parameters *ϕ* are calculated over minibatches of *m* training points, which are then used to update *ϕ* using the optimizer [28]. One training *epoch* is completed by iterating over all minibatches in the training dataset and hence performing many gradient steps (which can lead to faster convergence). Before training we initialize the network weights using the Glorot Uniform method [69].

The batch size and the learning rate are two optimizer hyperparameters that may significantly affect the results of the training procedure. We adjust these hyperparameters using heuristic arguments outlined below.

#### Batch Size

It has been noted that large batch size m may reduce the model’s ability to generalize, whereas small *m* can lead to more reliable results [70]. In our experiments we observed that very small batch sizes did little to improve results while significantly increasing training time. To balance these observations, we usually choose *m* to be 2-10% the size of the training dataset, which consistently gave good performance.

#### Learning Rate

The learning rate *η* of the optimizer controls the step size of each gradient update and should be chosen appropriately: too low a choice can lead to slow convergence, whereas a large learning rate can overshoot the target minimum. In our experiments we usually initialize *η* = 0.01 and decrease it as training progresses. Namely, we periodically monitor the loss function over the validation dataset and halve *η* if the loss has improved by less than 0.5% over the last 20 epochs on average; training terminates after *η* has been decreased 5 times, which usually indicates that optimization has stalled. We found this method to work well in our experiments.

This learning rate decay is similar to early stopping [71], a regularization technique that helps to prevent overfitting of the training dataset. Overfitting occurs when the neural network learns, or “memorizes”, particular features of the training dataset that are not representative of the model as a whole, and loses its ability to generalize to unseen data. This can often be detected by an increase in the validation loss together with a monotonically decreasing training loss. While a number of popular regularization strategies can be used to prevent this, such as L2 regularization or dropout [28], we did not find overfitting to be an issue in our experiments. We conjecture that this is due to the rigid nature of the negative binomial distribution which, unlike a Gaussian, cannot overfit single datapoints away from 0.

### B Hyperparameter Tuning

As noted in Section 2.2, training a neural network effectively requires finding a good architecture and hyperparameters. Beyond manual tuning this is classically done using black-box optimization methods such as grid search, random search or Bayesian Optimization [72]; the related [21] uses a more recent differentiable architecture search. For deep neural networks such approaches can be very computationally expensive depending on the number of hyperparameters. In contrast, our approach based on a single hidden layer can feasibly be tuned manually, and using the autoregulatory feedback loop in Section 4.1 as a testbed we obtain intuition about the effect of each hyperparameter for our problem.

#### Hidden Layers

The number and structure of the hidden layers in a neural network greatly affect its capacity, i.e. its ability to represent sufficiently complex functions. In Fig. 8**a** we plot the Hellinger distance between the true and predicted distributions for a single hidden layer with different numbers of neurons. We observe that the network’s capacity quickly grows with the number of neurons, reaching peak accuracy at about 128 neurons. Increasing the number further does not have a measurable effect on our model’s performance beyond increasing training time.

**Figure 8:**
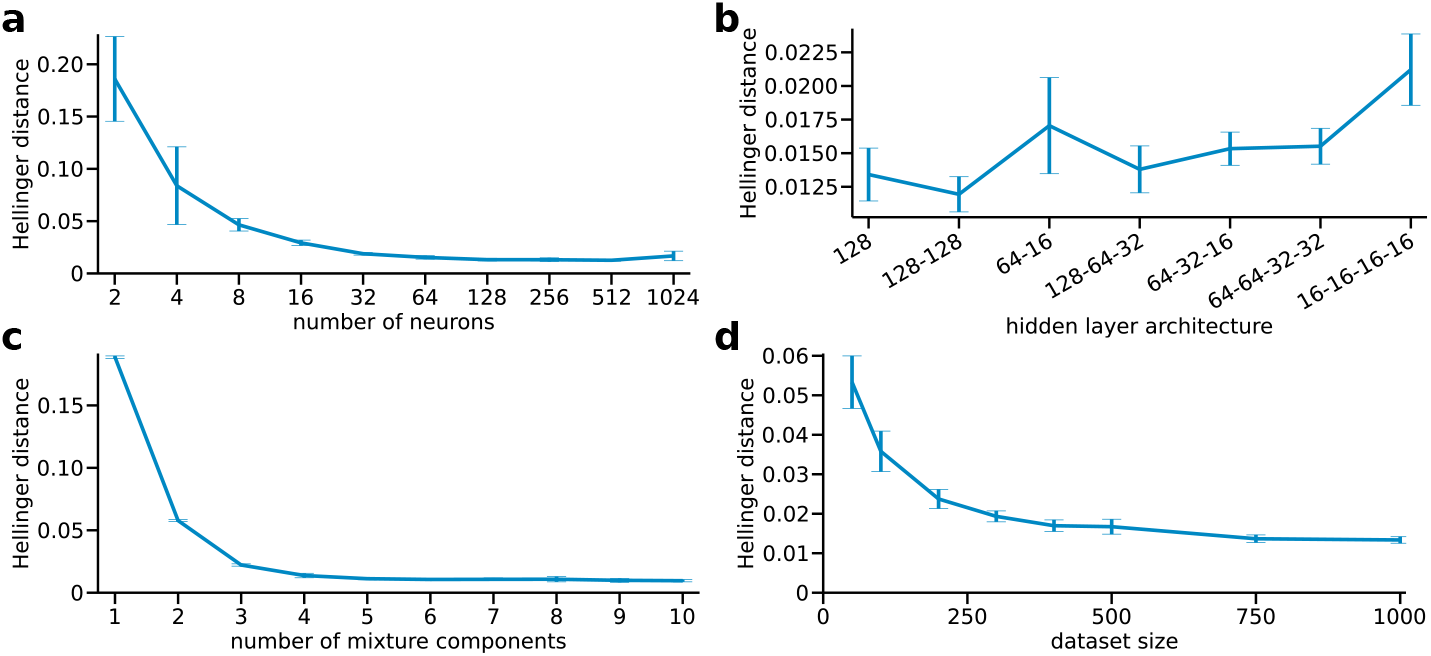
Results of manual hyperparameter tuning. Each point in the figure is the mean Hellinger distance for the validation dataset (consisting of 100 different parameter values at 4 time snapshots constructed with the FSP), computed over 10 independent training runs with a differently initialized neural network. For each run, we reinitialize the network, construct new training and validation datasets by drawing new parameters from a fixed Sobol sequence and train the network again. Note that the validation dataset used to produce the figure is distinct from the validation data used during training.

#### Network Depth

In Fig. 8**b** we compare the predictive performance for neural networks with multiple hidden layers of different sizes. The results suggest that shallow neural networks consisting of a single hidden layer are as effective as deeper ones for our purposes, while being easier to set up and train. Our experiments with the other systems (not shown) corroborated this observation.

#### Number of Components

Another important hyperparameter is the number of negative binomial components in the output mixture. In Fig. 8**c** we see that at least 4 components are needed to obtain good approximations for the autoregulatory feedback loop. This is not unexpected as the protein distributions of the autoregulatory feedback loop can be distinctly bimodal in certain parameter regimes, as shown in Fig. 3. Increasing the number of mixture components does not lead to overfitting, as observed in [26], but can increase the training time.

#### Dataset Size

Having enough training samples is essential to train a neural network in a way that allows it to learn to generalize. In Fig. 8**d** we show how increasing the size of the training dataset improves the performance of our network. Although relatively small datasets are sufficient for achieving good accuracy, in this case we have only five neural network inputs. Therefore, due to the curse of dimensionality, significantly larger datasets may be needed for effective training on chemical systems involving more reaction parameters. Adding more training points when the validation loss is significantly higher than the training loss is an effective way to determine the appropriate size.

#### Number of Simulations

The number of simulations at each training point also affects the accuracy of the fit. In general we suggest considering more SSA samples if the neural network does not provide a good fit on the training data. As fitting negative binomial mixtures to samples has a strong regularizing effect, the number of simulations per training point is generally much less than what is required to get an accurate histogram from samples (see Fig. 4).

### C Model Reaction Schemes

#### C.1 Autoregulatory Feedback Loop

The autoregulatory feedback loop presented in Section 4.1 is described by the following set of reactions (see also Fig. 3**a**):

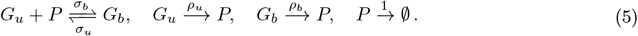

The specific parameter choices used for the analysis are indicated in Table 1.

#### C.2 Genetic Toggle Switch

The genetic toggle switch presented in Section 4.2 is modeled by the following reaction scheme (see also Fig. 5**a**):

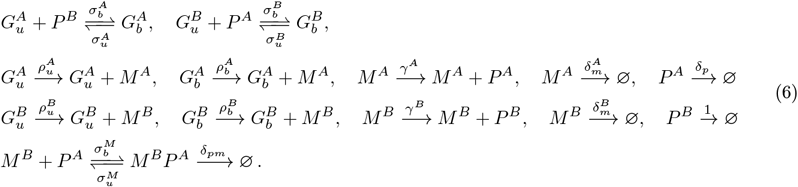

The specific parameter values used for the analysis are given in Table 2.

#### C.3 MAPK Pathway

The MAPK pathway presented in Section 4.3 is modeled by the following reaction scheme:

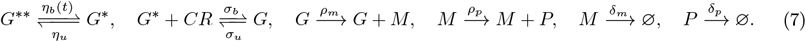

Here *η_b_*(*t*) depends on the current Hog1 concentration, which was measured experimentally, via the formula

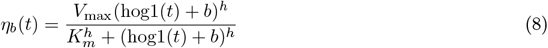

This is a simplification of the model presented in [56] assuming a fixed number of chromatin remodellers (*CR*) and ribosomes. The number of ribosomes does not change over time and was therefore absorbed into the protein production rate *ρ_p_*.

### D Model Parameters

